# Ion transport modulators differentially modulate inflammatory responses in THP-1 derived macrophages

**DOI:** 10.1101/2021.03.21.436302

**Authors:** Steven C. Mitini-Nkhoma, Narmada Fernando, G.K.D. Ishaka, Shiroma M. Handunnetti, Sisira L. Pathirana

## Abstract

Ion transport modulators are most commonly used to treat various non-communicable diseases including diabetes and hypertension. They are also known to bind to receptors on various immune cells, but the immunomodulatory properties of most ion transport modulators have not been fully elucidated. We assessed the effects of thirteen FDA approved ion transport modulators namely ambroxol HCl, amiloride HCl, diazoxide, digoxin, furosemide, hydrochlorothiazide, metformin, omeprazole, pantoprazole, phenytoin, verapamil, drug X and drug Y on superoxide production, nitric oxide production and cytokine expression by THP-1 derived macrophages that had been stimulated with ethanol-inactivated *Mycobacterium bovis* BCG. Ambroxol HCl, diazoxide, digoxin, furosemide, hydrochlorothiazide, metformin, pantoprazole, phenytoin, verapamil and drug Y had an inhibitory effect on nitric oxide production, while all the test drugs had an inhibitory effect on superoxide production. Amiloride HCl, diazoxide, digoxin, furosemide, phenytoin, verapamil, drug X and drug Y enhanced the expression of IL-1β and TNF-α. Unlike most immunomodulatory compounds currently in clinical use, most of the test drugs inhibited some inflammatory processes while promoting others. Ion pumps and ion channels could therefore serve as targets for more selective immunomodulatory agents which do not cause overt immunosupression.

## Introduction

The use of immunomodulators has increased significantly over the last few decades, in part due to a rise in the prevalence of autoimmune diseases worldwide [1,2]. Corticosteroids and nonsteroidal anti-inflammatory drugs (NSAIDS) are two of the oldest and most commonly used classes of immunomodulators in clinical practice. The term ‘corticosteroid’ encompasses various steroid hormones produced by the adrenal cortex and their synthetic analogues [3]. Corticosteroids bind to cytoplasmic steroid receptors, following which the receptor-ligand complex traverses the nuclear membrane and modulates transcription of various genes [3]. In addition to modulating transcription, corticosteroids can also directly modulate the activity of various proteins including G-protein coupled receptors [4]. Corticosteroids induce a wide range of physiological changes, and are thus associated with numerous adverse effects including osteoporosis and Cushing’s syndrome [5].

NSAIDS on the other hand have a relatively narrow activity spectrum, and primarily inhibit the activity of cyclooxygenase (COX) 1 and 2. COX1 and COX2 catalyse the production of prostaglandins, which mediate various inflammatory processes [6,7]. However, as prostaglandins are also involved in protection of the gastric mucosa from gastrointestinal secretions, NSAIDS at times cause peptic ulceration [8]. More selective COX2 inhibitors are generally less likely to cause peptic ulceration, but are associated with an increased risk of thrombosis [7,9].

As the demand for immunomodulators continues to increase, there is need for safer immunomodulatory agents. Over the last few decades there has been increasing interest in the use of ion transport modulators as immunomodulatory agents. Ion transport modulators are a diverse group of compounds that alter cell physiology by attuning ion currents across cellular and subcellular membranes. They are most commonly used to treat various non-communicable diseases including diabetes and hypertension. While most ion channel modulators directly interact with ion transporters and ion channels, others including metformin indirectly alter ion transport by interfering with processes upstream to activation of the ion transporters and channels [10]. As the immunomodulatory activities of most ion transport modulators have not been fully characterised, we conducted this study to assess the immunomodulatory properties of thirteen FDA approved ion transport modulators namely ambroxol HCl, amiloride HCl, diazoxide, digoxin, furosemide, hydrochlorothiazide (HCTZ), metformin, omeprazole, pantoprazole, phenytoin, verapamil, drug X and drug Y. We recently reported that these compounds also have potent antimycobacterial activity [11]. We have withheld the identities of drug X and drug Y pending further studies.

## Materials and Methods

### Macrophages

THP-1, a human monocytic cell line, was obtained from American Type Culture Collection (ATCC) and cultured in RPMI 1640 (catalog no. R6504; Sigma-Aldrich, St Louis, MO, USA) supplemented with 10% FBS (catalog no. 30-2020; ATCC), 10 mM HEPES (catalog no. H4034; Sigma-Aldrich), 4500 mg/l glucose (catalog no. G5500; Sigma-Aldrich), 1500 mg/l sodium bicarbonate (catalog no. S5761; Sigma-Aldrich) and 0.05 mM 2-mercaptoethanol (catalog no. M6250; Sigma-Aldrich). Prior to each experiment, the THP-1 cells were differentiated into macrophages by treatment with 200 nM phorbol-12-myristate-13-acetate (PMA) (catalog no. P8139; Sigma-Aldrich) for 3 days.

### *Mycobacterium bovis* BCG

*Mycobacterium bovis* BCG-1 (Russia) was kindly donated by Citihealth Imports (Pvt) limited. It was grown in BD Difco Middlebrook 7H9 broth (catalog no. DF0713-17-9, Thermo Fisher Scientific) supplemented with 10% (v/v) oleic acid-albumin-dextrose-catalase (OADC) growth supplement (catalog no. B12351, Thermo Fisher Scientific) and 0.05% tween 80 (catalog no. P4780; Sigma-Aldrich) to mid-log phase. The bacteria were then pelleted by centrifugation at 3000 *g* for 15 minutes. The supernatant was decanted and the pellet was resuspended in 70 % ethanol and kept at room temperature for 2 hours. The bacteria were then centrifuged again at 3000 *g* for 15 minutes, and the pellet was washed twice with PBS. The pellet was then re-suspended in PBS at a concentration of 3 × 10^6^ CFU/ml, aliquoted and stored at −20 °C until further use. To confirm that the *M. bovis* BCG had been inactivated, we plated it on BD Difco middlebrook 7H10 media (catalog no. DF0627-17-4; Thermo Fisher Scientific) and observed no growth.

### Drugs

All drugs that were tested in this study were purchased from Sigma-Aldrich. Stock solutions were prepared in dimethylsulfoxide (DMSO) (catalog no. C6295; Sigma-Aldrich), aliquoted, stored at −20 °C and used within six weeks. Working solutions were prepared in RPMI immediately before use. We assessed toxicity of the drugs using the Sulforhodamine B assay as previously described [12], and found that all the test drugs were not toxic to the macrophages at the concentrations that were used in this study (Table 1 below, and Figure S1 in Supplementary Materials). We used hydrocortisone (catalog no. H4001; Sigma-Aldrich), one of the most commonly used corticosteroids as a control in all the experiments, at a concentration of 75 ng/ml.

**Table 1:** Concentrations of drugs tested in the current study and their maximum free plasma concentrations in humans when taken at therapeutic doses.

| Drug | Concentration used in the current study in ng/ml | Maximum free plasma concentration in ng/ml |
| --- | --- | --- |
| Ambroxol HCl | 6 | 6 [13] |
| Amiloride HCl | 40 | 40 [14] |
| Diazoxide | 5000 | 5000 [15] |
| Digoxin | 2 | 2 [16] |
| Furosemide | 500 | 500 [17] |
| HCTZ | 500 | 642 [18] |
| Metformin | 1000 | 1000 [19] |
| Omeprazole | 200 | 200 [20] |
| Pantoprazole | 180 | 180 [21] |
| Phenytoin | 2000 | 2000 [22] |
| Verapamil | 24 | 24 [23] |
| Drug X | 2 | 2 |
| Drug Y | 2 | 2 |

Note. Reprinted from "Ion transport modulators as antimycobacterial agents," by SC Mitini-Nkhoma et al, 2020, Tuberculosis Research and Treatment, vol. 2020, https://doi.org/10.1155/2020/3767915.

### Assessment of effects of ion transport modulators on nitric oxide production

Nitric oxide (NO) is converted into nitrites (NO_2_^−^) and nitrates (NO_3_^−^) seconds after it is synthesized. NO_2_^−^ and NO_3_ ^−^ are therefore often used as surrogate measures for NO production [24]. We assessed total NO_x_ (NO_2_^−^ + NO_3_^−^) levels using the modified Griess assay as previously described, with slight modifications [25]. Griess reagent was prepared in-house by mixing equal volumes of 0.1% N-(naphtyl)ethelenediamine (NED) (catalog no. 222488; Sigma-Aldrich) in distilled water with 1% Sulfanilamide (catalog no. 46874; Sigma-Aldrich) in 5% phosphoric acid (catalog no. W290017; Sigma-Aldrich) just before use.

We seeded 2 × 10^4^ macrophages into each well of a 96 well plate, and stimulated them with 2 × 10^5^ CFU of ethanol-inactivated *M. bovis* BCG, in the presence or absence of each of the test drugs for 24 hours at 37 °C. The supernatant was then harvested and centrifuged at 5000 *g* for 10 minutes. Equal volumes of supernatant, Griess reagent and vanadium (III) chloride (catalog no. 208272; Sigma-Aldrich) were mixed together and incubated for 30 minutes. We then measured the absorbance of the resulting azo dye at 540 nm using a multi-mode microplate reader (Synergy™ HTX, Biotek, USA). The amount of NO_x_ produced in each culture was then interpolated from a standard curve generated using sodium nitrite.

### Assessment of the effect of ion transport modulators on superoxide production

Superoxide production was assessed using the nitroblue tetrazolium (NBT) assay as previously described [26]. Superoxide reduces NBT to NBT-formazan, which can then be quantified spectrophotometrically to estimate the rate of superoxide production. We seeded 2 × 10^4^ macrophages into each well of a 96 well plate in RPMI, and stimulated them with 2 × 10^5^ CFU of ethanol-inactivated *M. bovis* BCG in the presence of NBT (catalog no N6876, Sigma-Aldrich) and the test drugs over 30 minutes. The supernatant was then discarded and the macrophages were washed twice with PBS. The macrophages, which now contained NBT-formazan, were fixed with 70% methanol (catalog no. 179337, Sigma-Aldrich) and allowed to air dry. The NBT-formazan was solubilized in dissolving media containing dimethyl sulfoxide and potassium hydroxide (catalog no P5958, Sigma-Aldrich). The suspension was centrifuged to remove cell debris, after which the amount of NBT-formazan in the supernatant was quantified by measuring absorbance at 620 nm using a multi-mode microplate reader. The amount of NBT reduced in the drug-treated cultures was calculated as a percentage of that reduced by the drug-free controls as follows:

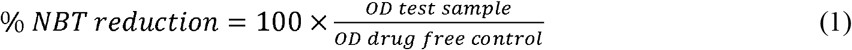

### Assessment of the effects of ion transport modulators on expression of cytokine genes

Expression of IL-1β, IL-10, IL-12α and TNF-α genes was assessed by RT-qPCR. We seeded 1 × 10^6^ macrophages into each well of a 6 well plate, and stimulated them with 10 × 10^6^ CFU of ethanol-inactivated *M. bovis* BCG in the presence or absence of the test drugs for 24 hours. RNA was then extracted using the RNEasy mini kit (Qiagen, Hilden, Germany), after which cDNA was synthesised using the GoScript reverse transcription system (Promega, Madison, USA). Quantitative PCR was performed using the Mesa green qPCR master mix (Eurogentec, Seraing, Germany). All primer sequences were obtained from Primerbank (Table 2), and the primers were synthesized by Integrated DNA Technologies (IDT).

**Table 2:**
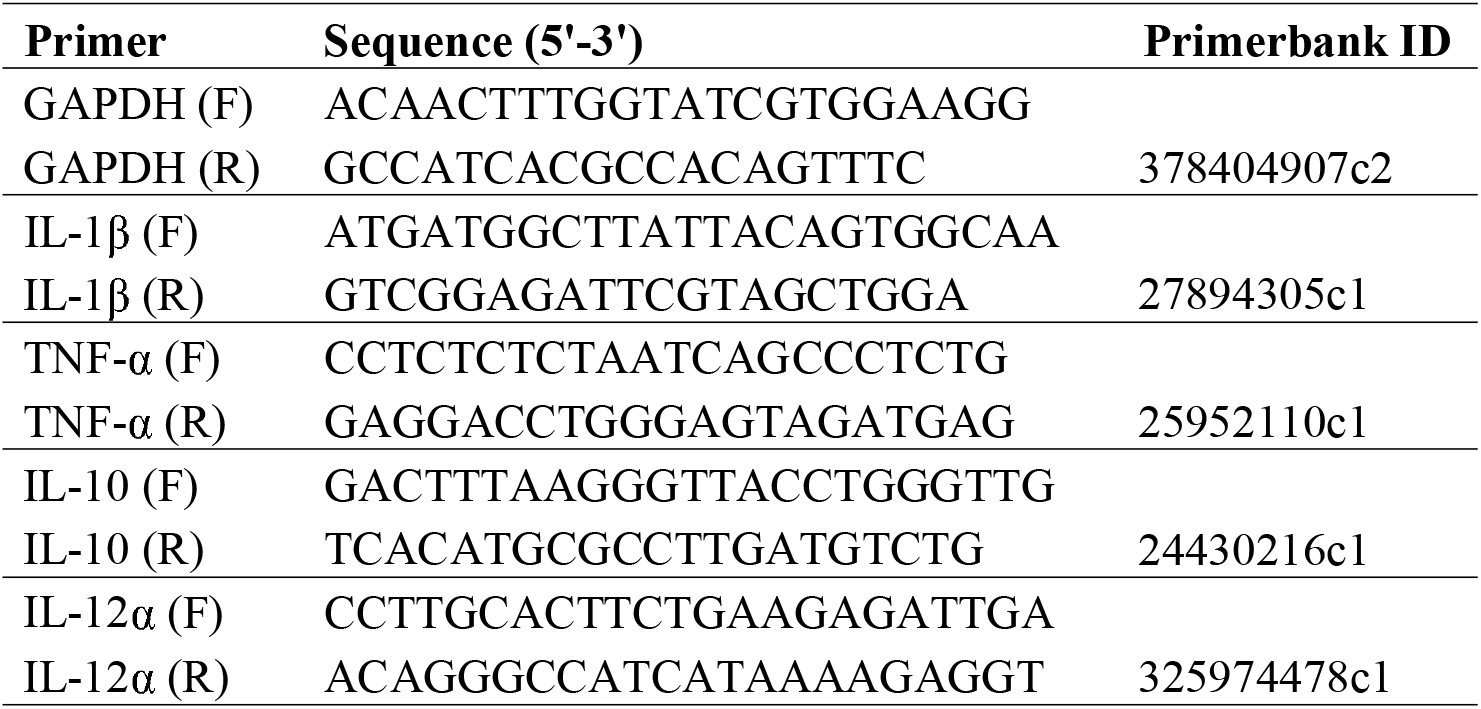
Primers used for quantification of cytokine RNA.

### Data analysis

Data from the NBT and Griess assays were analysed using GraphPad Prism version 8.4.2 (GraphPad Software, La Jolla, California, USA). Production of nitric oxide and superoxide in the drug-treated samples and drug-free controls was compared using one-way analysis of variance (ANOVA) followed by Dunnett’s post hoc test.

RNA expression data was analysed using R version 3.5.1 and the pcr library [27,28]. We used the Livak method to estimate expression of the genes in the treated cultures relative to the drug-free controls [29], with GAPDH as a reference gene.

## Results and discussion

### Ion transport modulators alter nitric oxide production in macrophages

Nitric oxide production (indicated by total NO_x_ levels) was lower in macrophages that were treated with ambroxol HCl, diazoxide, digoxin, furosemide, HCTZ, metformin, pantoprazole, phenytoin, verapamil or drug Y than in drug-free controls (p<0.001, Figure 1). Macrophages that were treated with amiloride HCl produced more nitric oxide than drug-free controls (p<0.001). There was no significant difference in nitric oxide production between drug-free controls and cultures that were treated with omeprazole or drug X.

Our findings are consistent with those of various authors, including Shen et al. (1995), who demonstrated that verapamil inhibits nitric oxide production in PMA-stimulated mouse peritoneal macrophages [30]. Verapamil inhibits the entry of calcium into cells through L-type voltage-gated calcium channels (VGCCs). Calcium is an important second messenger in several signal transduction pathways involved in macrophage activation [31]. Verapamil might therefore modulate nitric oxide production by interfering with calcium signalling.

**Figure 1:**
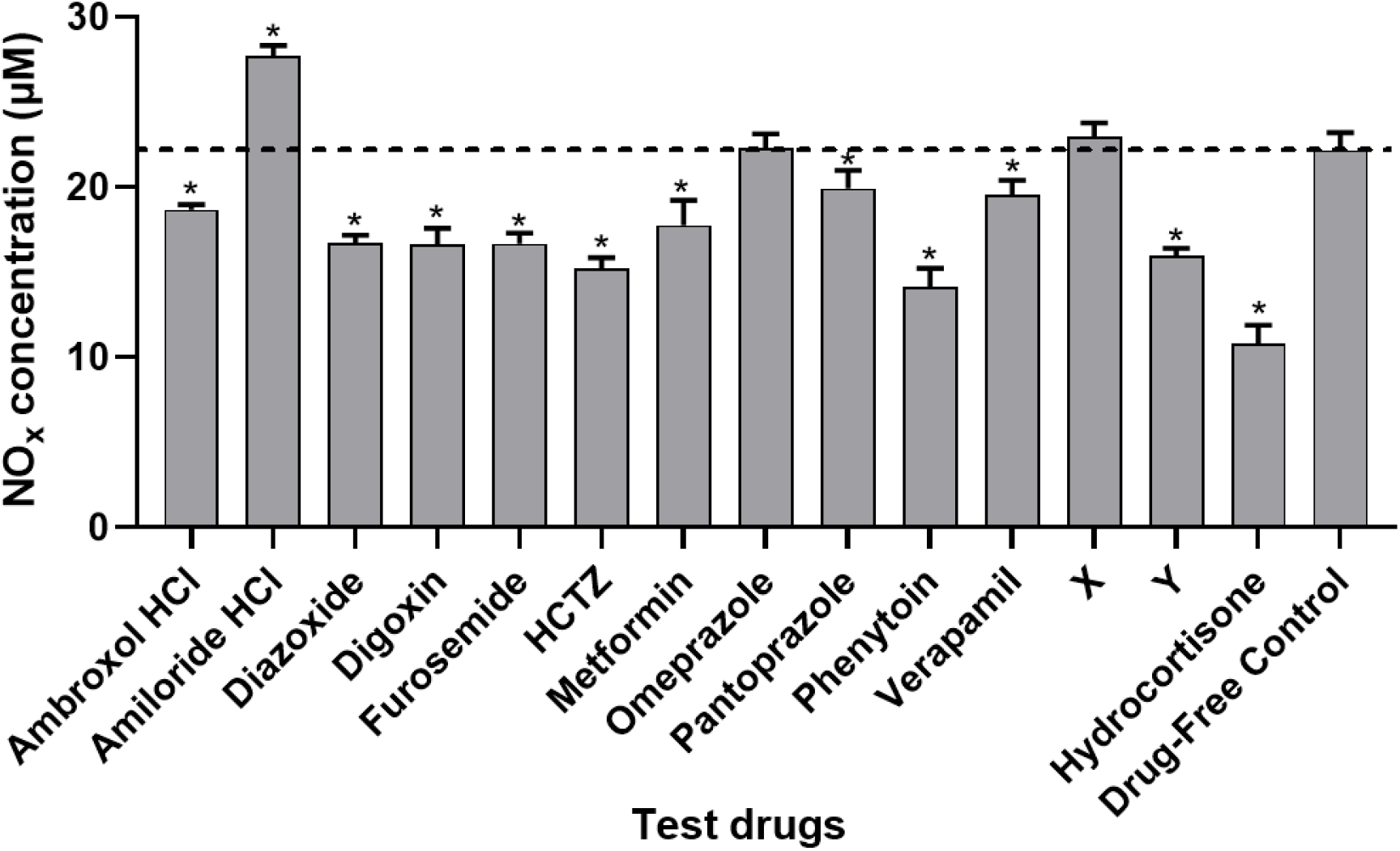
Effects of ion transport modulators on production of nitrites and nitrates by THP-1 derived macrophages. Macrophages were stimulated with ethanol-inactivated *M. bovis* BCG in the presence or absence of the test drugs over 24 hours. Nitric oxide production was then assessed by quantifying NO_x_ levels in the supernatant using the Griess assay. Nitric oxide production was compared between the different cultures using one way analysis of variance (ANOVA) followed by Dunnett’s post hoc test. Data are presented as mean + SD of six replicates pooled from two independent experiments. Dashed line indicates nitric oxide production in drug-free controls. * indicates p<0.05 vs drug-free controls. HCTZ, hydrochlorothiazide.

Phenytoin and ambroxol HCl are potent inhibitors of voltage-gated sodium channels (Na_V_s) 1.5 and 1.8 respectively [32,33]. Various authors have previously documented the inhibitory effects of both Na_V_ inhibitors on nitric oxide production [34–36]. Phenytoin also inhibits various other macrophage functions including metalloproteinase production, which contributes to gingival hypertrophy in some people on chronic phenytoin therapy [37]. While the physiology of Na_V_ 1.5 has been most extensively studied in cardiac and neural tissue, Parpalardo et al. (2014) demonstrated that knockdown of Na_V_ 1.5 attenuates calcium influx in astrocytes [38]. Blocking Na_V_s might therefore indirectly modulate leukocyte function by modulating calcium signalling.

Our findings also echo those of Kato et al. (2010), who demonstrated that metformin inhibits nitric oxide production in lipopolysaccharide (LPS)-stimulated macrophages [39]. Interestingly, some studies have demonstrated that metformin enhances nitric oxide production, particularly in the absence of a pro-inflammatory stimuli [40–42]. Nitric oxide production is mediated by 3 isoforms of the enzyme nitric oxide synthase (NOS); namely neuronal NOS (nNOS), inducible NOS (iNOS) and endothelial NOS (eNOS) [43]. Of the three, nNOS and eNOS are generally expressed constitutively in various cell types, while expression of iNOS is induced by pro-inflammatory stimuli [43]. It is therefore likely that metformin has opposing effects on different isoforms of NOS.

Our results also indicate that diazoxide inhibits nitric oxide production in macrophages. This is in agreement with the findings of Virgili et al (2011), who demonstrated that diazoxide inhibits production of nitric oxide in murine model of multiple sclerosis [44]. Diazoxide is an agonist of ATP-gated potassium channels primarily used to treat patients with hypoglycaemia [15]. While calcium is more abundant in the extracellular fluid, potassium is highly concentrated in the cytosol. Opening of plasma membrane potassium channels leads to efflux of potassium down its chemical gradient, thus increasing the electrical gradient between the cytosol and the extracellular fluid. This increases the driving force for calcium influx. Diazoxide might therefore modulate nitric oxide production by indirectly modulating calcium signalling.

In the present study, macrophages that were treated with amiloride HCl produced more nitric oxide than drug-free controls. Amiloride HCl is primarily considered an inhibitor of Na+/H+ exchangers (NHEs), but is also known to inhibit the activity of various sodium and calcium channels [45]. There is therefore need for further studies to determine how amiloride HCl promotes nitric oxide production. While nitric oxide contributes to the pathogenesis of various diseases, it ameliorates pathology in others. Inhailed nitric oxide is used to treat various cardiopulmonary illnesses including acute respiratory distress syndrome [46]. Amiloride HCl and other agents which enhance nitric oxide production could potentially provide a cheaper and more convenient alternative to inhaled nitric oxide. Conversely, ambroxol HCl, diazoxide, digoxin, furosemide, HCTZ, metformin, pantoprazole, phenytoin, verapamil and drug Y, which demonstrated an inhibitory effect on production of nitric oxide in this study, could potentially be used to ameliorate pathology in patients with septic shock and other diseases that involve anomalous production of nitric oxide.

### Ion transport modulators alter superoxide production by macrophages

Superoxide production (indicated by % NBT reduction) was lower in macrophages that were treated with any of the test drugs than in the drug-free controls (Figure 2). Macrophages that were treated with verapamil produced the least amount of superoxide (62.4%, p<0.001). Our findings are in line with previous findings which have demonstrated that amiloride HCl, ambroxol HCl, metformin, omeprazole and verapamil inhibit superoxide production by leukocytes [47–51]. Inhibition of superoxide production by the proton pump inhibitors (PPIs) omeprazole and pantoprazole, which are most commonly used to treat peptic ulcer disease and gastritis, might be one of the reasons for the high prevalence of gastrointestinal bacterial infections in people on chronic PPI therapy.

**Figure 2:**
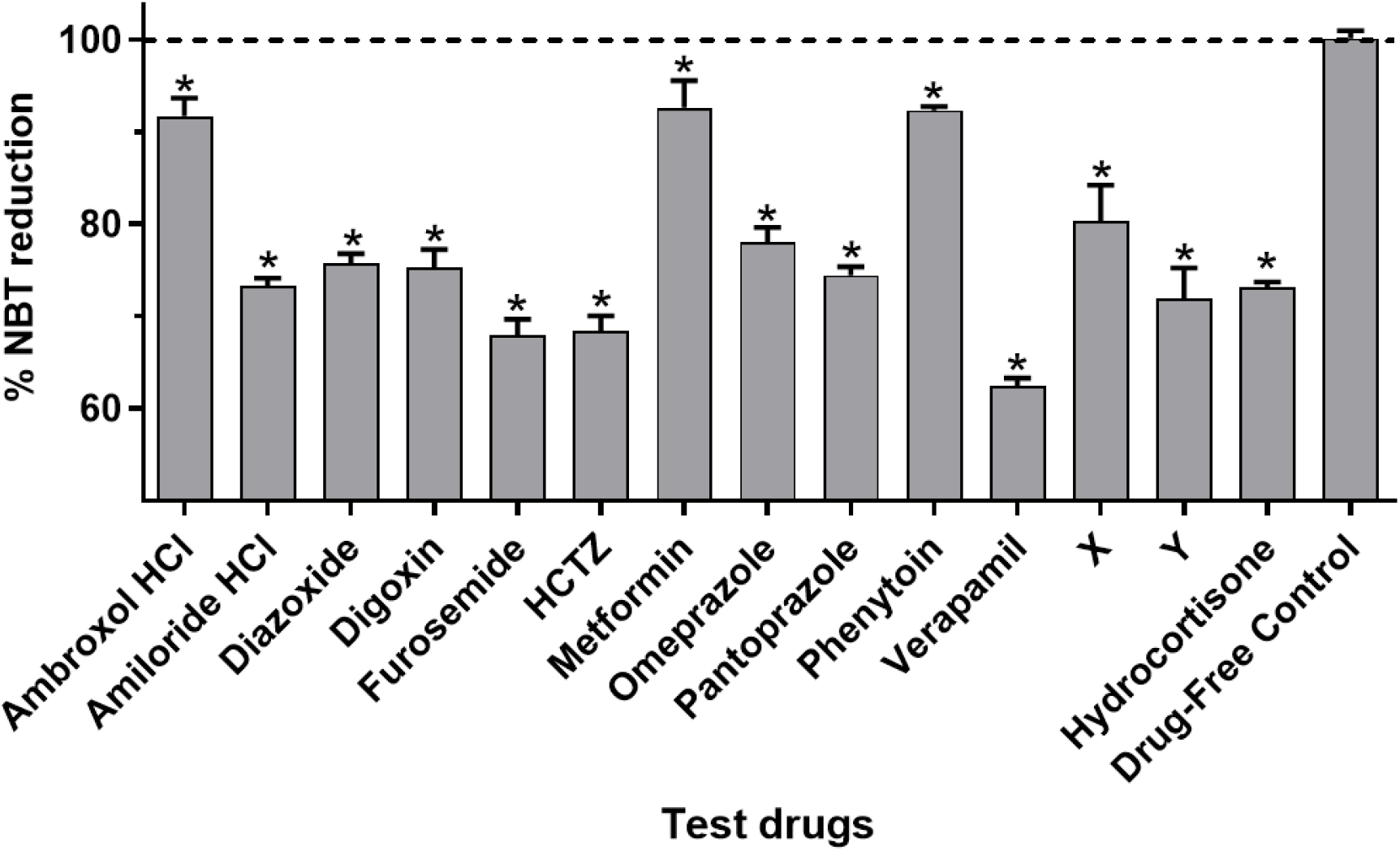
Effects of ion transport modulators on production of superoxide by THP-1 derived macrophages. Macrophages were stimulated with ethanol-inactivated *M. bovis* BCG in the presence of the test drugs over 30 minutes, and production of superoxide was assessed using the NBT assay. Superoxide production was compared between the different cultures using one way analysis of variance (ANOVA) followed by a Dunnett’s post hoc test. Data are presented as mean (+SD) of six replicates pooled from two independent experiments. Dotted line indicates superoxide production in drug-free controls. * indicates p<0.05 vs drug-free controls. HCTZ, hydrochlorothiazide.

In macrophages, NADPH oxidases are the key producers of superoxide [52]. Shen et al. (1995) demonstrated that the calcium signalling pathway is involved in the activation of the NADPH oxidase [30]. As was the case with nitric oxide production, most of the compounds might have therefore inhibited superoxide production by indirectly modulating calcium signalling. In addition to NADPH oxidases, the three isoforms of NOS also contribute to superoxide production in macrophages [53]. Therefore, the test drugs also might have reduced nitric oxide production in the macrophages by inhibiting the activity of iNOS.

Hydrocortisone is one of the most commonly used corticosteroids in clinical practice. In the present study, superoxide production was lower in macrophages that were treated with furosemide, HCTZ, verapamil or drug Y, than macrophages that were treated with hydrocortisone. Unlike ion transport modulators, hydrocortisone primarily functions by altering cellular transcription patterns, and thus has a slow onset [3]. The effect of hydrocortisone on macrophages was therefore likely not maximal at the time the experiment was completed.

As superoxide is implicated in the pathogenesis of various diseases including atherosclerosis, myocardial infarction and cerebrovascular accidents [54], ion channel modulators could potentially be used to ameliorate pathology in such diseases.

### Ion transport modulators alter expression of cytokines by macrophages

Macrophages that were treated with amiloride HCl, diazoxide, digoxin, furosemide, phenytoin, verapamil drug X and drug Y had higher levels of expression of TNF-α and IL-1β genes than drug-free controls (p<0.001, Figure 3). Macrophages that were treated with metformin and pantoprazole had higher levels of expression of TNF-α but not IL-1β. Expression of TNF-α was highest in macrophages that were treated with furosemide, while that of IL-1β was highest in macrophages that were treated with phenytoin. There was no statistically significant difference in expression of TNF-α and IL-1β between cultures that were treated with ambroxol HCl or omeprazole and the drug-free controls.

**Figure 3:**
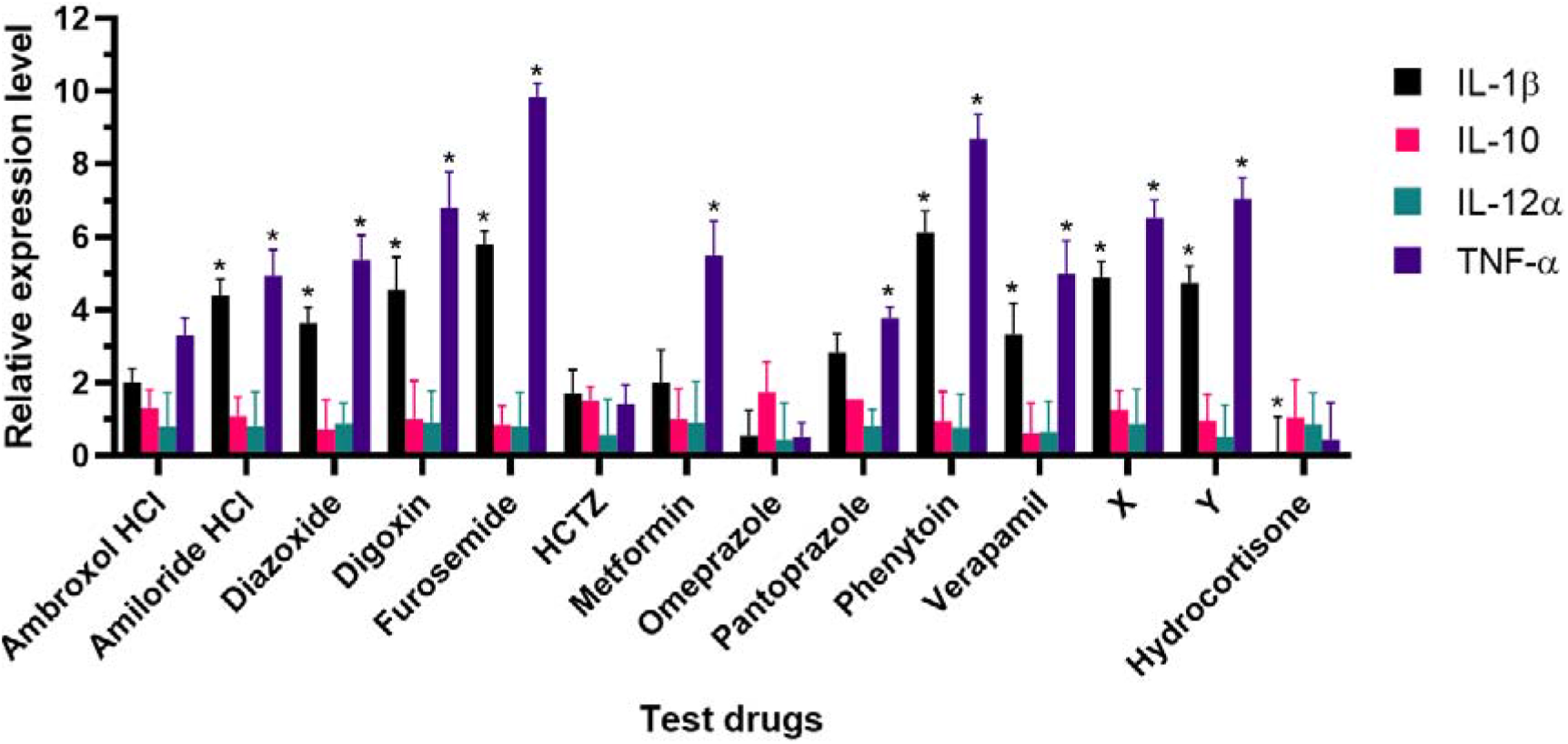
Effect of ion transport modulators on cytokine expression by macrophages. Macrophages were stimulated with ethanol-inactivated *M. bovis* BCG in the presence of the test drugs over 24 hours. RNA expression levels in the drug-treated cultures relative to the drug-free controls were calculated using the Livak method. Data are presented as mean (+SD) of two replicates. * indicates p<0.05 vs drug-free controls. HCTZ, hydrochlorothiazide.

We did not observe any statistically significant differences in the expression of IL-12α and IL-10 between the drug-free controls and any of the test conditions.

Our findings are in line with those of Gupta et al. (2009) who demonstrated that inhibiting the activity of L-type voltage gated calcium channels, enhances the expression of pro-inflammatory cytokines in macrophages and dendritic cells stimulated with *Mycobacterium tuberculosis* (Mtb) lysate [55]. However, Li et al. (2006) observed contrasting findings following concomitant exposure of Sprague-Dawley rats to LPS and verapamil [56]. The incongruity is likely because different stimulants invoke different combinations of cell activation pathways, leading to differences in cellular responses to both the stimulants and any concomitantly administered compounds. In addition, different cell types may respond differently to both the stimulants and the ion transport modulators.

As with verapamil, there is contrasting literature on the effects of the sodium channel agonists phenytoin and ambroxol HCl, and indeed other ion channel modulators on cytokine production by macrophages. For example, while Song et al. (1997) demonstrated that phenytoin promotes production of TNF-α by wound macrophages following systemic and local irradiation in mice[57], Jackson et al. (2019) observed that phenytoin inhibits TNF-α production in human multipotent adult progenitor cells [58]. In addition, Ichiyama et al. (2000) reported that phenytoin has no effect on production of TNF-α in LPS-stimulated THP-1 cells. The incongruency in the results obtained from different systems therefore indicates that ion transport modulators do not cause a global depression of inflammatory processes, and may enhance others depending on the context.

### Relationship between nitric oxide production, superoxide production and cytokine expression

Table 3 below summarises the immunomodulatory activities of the ion transport modulators tested in this study.

**Table 3:** Summary of immunomodulatory activity of ion transport modulators.

| | Nitric oxide | Superoxide | TNF-α | IL-1β | IL-10 | IL-12α |
| --- | --- | --- | --- | --- | --- | --- |
| Ambroxol HCl | ↓ | ↓ | ↔ | ↔ | ↔ | ↔ |
| Amiloride HCl | ↑ | ↓ | ↑ | ↑ | ↔ | ↔ |
| Diazoxide | ↓ | ↓ | ↑ | ↑ | ↔ | ↔ |
| Digoxin | ↓ | ↓ | ↑ | ↑ | ↔ | ↔ |
| Furosemide | ↓ | ↓ | ↑ | ↑ | ↔ | ↔ |
| HCTZ | ↓ | ↓ | ↔ | ↔ | ↔ | ↔ |
| Metformin | ↓ | ↓ | ↑ | ↔ | ↔ | ↔ |
| Omeprazole | ↔ | ↓ | ↔ | ↔ | ↔ | ↔ |
| Pantoprazole | ↓ | ↓ | ↑ | ↔ | ↔ | ↔ |
| Phenytoin | ↓ | ↓ | ↑ | ↑ | ↔ | ↔ |
| Verapamil | ↓ | ↓ | ↑ | ↑ | ↔ | ↔ |
| X | ↔ | ↓ | ↑ | ↑ | ↔ | ↔ |
| Y | ↓ | ↓ | ↑ | ↑ | ↔ | ↔ |

We pooled together the data obtained with all the test drugs in all the assays to assess the correlation between the different aspects of macrophage function (Figure 4). We observed a strong positive correlation between TNF-α and IL-1β expression (r=0.92, p=<0.001, Figure 4), and a strong negative correlation between the expression of IL-10 and TNF-α (r=-0.7, p=0.007). We also observed a negative albeit weak correlation between the production of nitric oxide and the expression of TNF-α (r=-0.31, p=0.308). This finding is consistent with that of Thomassen et al. (1998) who demonstrated that nitric oxide down-regulates the production of IL-1β and TNF-α in human alveolar macrophage [59]. Therefore, the test compounds might have enhanced the expression of TNF-α and IL-1β by inhibiting nitric oxide production, thus removing the tonic inhibition on the expression of pro-inflammatory cytokines.

**Figure 4:**
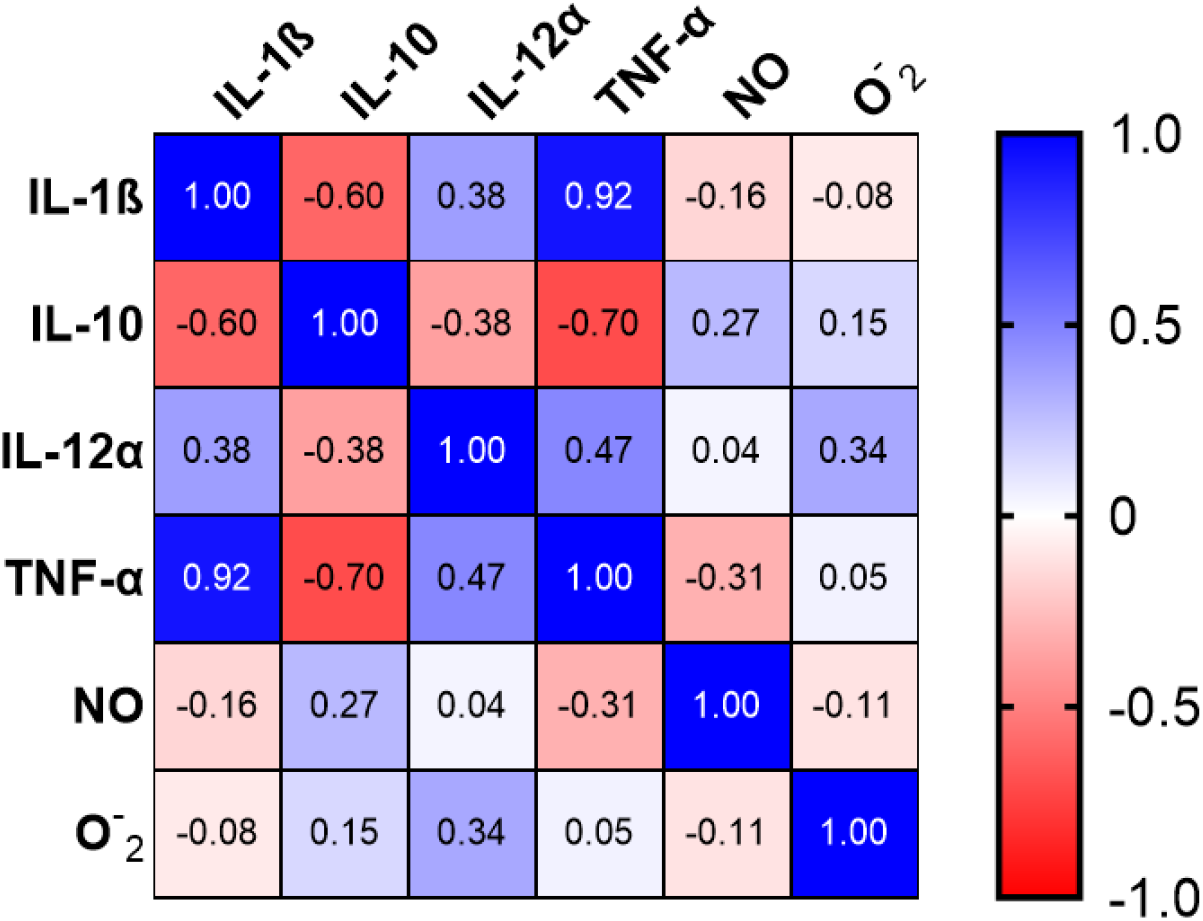
Spearman correlation coefficients between different markers of macrophage function.

## Conclusions

In summary, we have demonstrated that ambroxol HCl, amiloride HCl, diazoxide, digoxin, furosemide, HCTZ, metformin, omeprazole, pantoprazole, phenytoin, verapamil, drug X and drug Y have potent immunomodulatory properties. More importantly, the results indicate that the ion transport modulators differentially alter various aspects of macrophage function, unlike most immunomodulators in clinical practice which depress all immune responses. Therefore, ion transport modulators could potentially serve as more selective immunomodulators, with a lower risk of causing overt immunosuppression.

## Limitations

While we assessed the expression of cytokine genes, we could not assess their secretion. In addition, as this study was conducted *in vitro*, it might not accurately reflect how the ion channel modulators would affect macrophage physiology *in vivo*. There is therefore need to ascertain the immunomodulatory properties of the ion transport modulators tested in this study in *in vivo* systems.

## Data Availability

The data used to support the findings of this study are available from the authors upon request.

## Supplementary Materials

Figure S1: Mean (+SD) viability of THP-1 derived macrophages following exposure to different concentrations of test drugs.

## References

[1] G.S. Cooper, M.L.K. Bynum, E.C. Somers, Recent insights in the epidemiology of autoimmune diseases: Improved prevalence estimates and understanding of clustering of diseases, J. Autoimmun. 33 (2009) 197–207. https://doi.org/10.1016/j.jaut.2009.09.008.

[2] C.W. Schmidt, Questions persist: environmental factors in autoimmune disease., Environ. Health Perspect. 119 (2011) A248. https://doi.org/10.1289/ehp.119-a248.

[3] S. Ramamoorthy, J.A. Cidlowski, Corticosteroids. Mechanisms of Action in Health and Disease, Rheum. Dis. Clin. North Am. 42 (2016) 15–31. https://doi.org/10.1016/j.rdc.2015.08.002.

[4] C. Wang, Y. Liu, J.M. Cao, G protein-coupled receptors: Extranuclear mediators for the non-genomic actions of steroids, Int. J. Mol. Sci. 15 (2014) 15412–15425. https://doi.org/10.3390/ijms150915412.

[5] W. Ericson-Neilsen, A.D. Kaye, Steroids: Pharmacology, complications, and practice Delivery Issues, Ochsner J. 14 (2014) 203–207.

[6] J.R. Vane, Inhibition of prostaglandin synthesis as a mechanism of action for aspirin-like drugs, Nat. New Biol. 231 (1971) 232–235. https://doi.org/10.1038/newbio231232a0.

[7] A. Zarghi, S. Arfaei, Selective COX-2 inhibitors: A review of their structure-activity relationships, Iran. J. Pharm. Res. 10 (2011) 655–683. https://doi.org/10.22037/ijpr.2011.1047.

[8] S. Wongrakpanich, A. Wongrakpanich, K. Melhado, J. Rangaswami, A comprehensive review of non-steroidal anti-inflammatory drug use in the elderly, Aging Dis. 9 (2018) 143–150. https://doi.org/10.14336/AD.2017.0306.

[9] R.J. Bing, M. Lomnicka, Why do cyclo-oxygenase-2 inhibitors cause cardiovascular events?, J. Am. Coll. Cardiol. 39 (2002) 521–522. https://doi.org/10.1016/S0735-1097(01)01749-1.

[10] X. Fu, Y. Pan, Q. Cao, B. Li, S. Wang, H. Du, N. Duan, X. Li, Metformin restores electrophysiology of small conductance calcium-activated potassium channels in the atrium of GK diabetic rats, BMC Cardiovasc. Disord. 18 (2018) 63. https://doi.org/10.1186/s12872-018-0805-5.

[11] S.C. Mitini-Nkhoma, N. Fernando, G.K.D. Ishaka, S.M. Handunnetti, S.L. Pathirana, Ion Transport Modulators as Antimycobacterial Agents, Tuberc. Res. Treat. 2020 (2020) 1–7. https://doi.org/10.1155/2020/3767915.

[12] S.R. Samarakoon, I. Thabrew, P. Galhena, D. De Silva, K. Tennekoon, A comparison of the cytotoxic potential of standardized aqueous and ethanolic extracts of a polyherbal mixture comprised of Nigella sativa (seeds), Hemidesmus indicus (roots) and Smilax glabra (rhizome), Pharmacognosy Res. 2 (2010) 335–342. https://doi.org/10.4103/0974-8490.75451.

[13] Y.-G. Yang, L.-X. Song, N. Jiang, X.-T. Xu, X.-H. Di, M. Zhang, Pharmacokinetics of ambroxol and clenbuterol tablets in healthy Chinese volunteers., Int. J. Clin. Exp. Med. 8 (2015) 18744–50.

[14] K.M. Jones, E. Liao, K. Hohneker, S. Turpin, M.M. Henry, K. Selinger, P.H. Hsyu, R.C. Boucher, M.R. Knowles, G.E. Dukes, Pharmacokinetics of amiloride after inhalation and oral administration in adolescents and adults with cystic fibrosis., Pharmacotherapy. 17 (n.d.) 263–70.

[15] R. Kizu, K. Nishimura, R. Sato, K. Kosaki, T. Tanaka, Y. Tanigawara, T. Hasegawa, Population Pharmacokinetics of Diazoxide in Children with Hyperinsulinemic Hypoglycemia., Horm. Res. Paediatr. 88 (2017) 316–323. https://doi.org/10.1159/000478696.

[16] T.W. Smith, V.P. Butler, E. Haber, Determination of Therapeutic and Toxic Serum Digoxin Concentrations by Radioimmunoassay, N. Engl. J. Med. 281 (1969) 1212–1216. https://doi.org/10.1056/NEJM196911272812203.

[17] University of Lausanne, Furosemide, (2018). https://sepia2.unil.ch/pharmacology/index.php?id=102 (accessed June 7, 2019).

[18] R. Aubin, P. Ménard, D. Lajeunesse, Selective effect of thiazides on the human osteoblast-like cell line MG-63, Kidney Int. 50 (1996) 1476–1482. https://doi.org/10.1038/ki.1996.461.

[19] F. Kajbaf, M.E. De Broe, J.-D. Lalau, Therapeutic Concentrations of Metformin: A Systematic Review, Clin. Pharmacokinet. 55 (2016) 439–459. https://doi.org/10.1007/s40262-015-0323-x.

[20] J.M. Shin, N. Kim, Pharmacokinetics and pharmacodynamics of the proton pump inhibitors, J. Neurogastroenterol. Motil. 19 (2013) 25–35. https://doi.org/10.5056/jnm.2013.19.1.25.

[21] R. Huber, M. Hartmann, H. Bliesath, R. Lühmann, V.W. Steinijans, K. Zech, Pharmacokinetics of pantoprazole in man., Int. J. Clin. Pharmacol. Ther. 34 (1996) S7–16.

[22] J. Galjour, Phenytoin Level, Medscape. (2014). https://emedicine.medscape.com/article/2090306-overview#a1 (accessed June 7, 2019).

[23] W. Frishman, E. Kirsten, M. Klein, M. Pine, S.M. Johnson, L.D. Hillis, M. Packer, R. Kates, Clinical relevance of verapamil plasma levels in stable angina pectoris, Am. J. Cardiol. 50 (1982) 1180–1184. https://doi.org/10.1016/0002-9149(82)90440-4.

[24] N.S. Bryan, M.B. Grisham, Methods to detect nitric oxide and its metabolites in biological samples, Free Radic. Biol. Med. 43 (2007) 645–657. https://doi.org/10.1016/j.freeradbiomed.2007.04.026.

[25] G.C. Park, J.S. Ryu, D.Y. Min, The role of nitric oxide as an effector of macrophage-mediated cytotoxicity against Trichomonas vaginalis., Korean J. Parasitol. 35 (1997) 189–195. https://doi.org/10.3347/kjp.1997.35.3.189.

[26] H.S. Choi, J.W. Kim, Y.-N. Cha, C. Kim, A quantitative nitroblue tetrazolium assay for determining intracellular superoxide anion production in phagocytic cells., J. Immunoassay Immunochem. 27 (2006) 31–44. https://doi.org/10.1080/15321810500403722.

[27] R Core Team, R: A Language and Environment for Statistical Computing, (2018).

[28] M. Ahmed, D.R. Kim, pcr: An R package for quality assessment, analysis and testing of qPCR data, PeerJ. 2018 (2018) e4473. https://doi.org/10.7717/peerj.4473.

[29] K.J. Livak, T.D. Schmittgen, Analysis of relative gene expression data using real-time quantitative PCR and the 2-ΔΔCT method, Methods. 25 (2001) 402–408. https://doi.org/10.1006/meth.2001.1262.

[30] H. Shen, M.D. Wiederhold, D.W. Ou, The suppression of macrophage secretion by calcium blockers and adenosine, Immunopharmacol. Immunotoxicol. 17 (1995) 301–309. https://doi.org/10.3109/08923979509019752.

[31] A.C. Newton, M.D. Bootman, J. Scott, Second messengers, Cold Spring Harb. Perspect. Biol. 8 (2016). https://doi.org/10.1101/cshperspect.a005926.

[32] W. Gaida, K. Klinder, K. Arndt, T. Weiser, Ambroxol, a Nav1.8-preferring Na+ channel blocker, effectively suppresses pain symptoms in animal models of chronic, neuropathic and inflammatory pain, Neuropharmacology. 49 (2005) 1220–1227. https://doi.org/10.1016/j.neuropharm.2005.08.004.

[33] M. Yang, D.J. Kozminski, L.A. Wold, R. Modak, J.D. Calhoun, L.L. Isom, W.J. Brackenbury, Therapeutic potential for phenytoin: Targeting Nav1.5 sodium channels to reduce migration and invasion in metastatic breast cancer, Breast Cancer Res. Treat. 134 (2012) 603–615. https://doi.org/10.1007/s10549-012-2102-9.

[34] L.P. Reagan, C.R. McKittrick, B.S. McEwen, Corticosterone and phenytoin reduce neuronal nitric oxide synthase messenger RNA expression in rat hippocampus, Neuroscience. 91 (1999) 211–219. https://doi.org/10.1016/S0306-4522(98)00615-0.

[35] Y.Y. Jang, J.H. Song, Y.K. Shin, E.S. Han, C.S. Lee, Depressant effects of ambroxol and erdosteine on cytokine synthesis, granule enzyme release, and free radical production in rat alveolar macrophages activated by lipopolysaccharide, Pharmacol. Toxicol. 92 (2003) 173–179. https://doi.org/10.1034/j.1600-0773.2003.920407.x.

[36] F.L.M. Ricciardolo, V. Sorbello, S. Benedetto, D. Paleari, Effect of Ambroxol and Beclomethasone on Lipopolysaccharide-Induced Nitrosative Stress in Bronchial Epithelial Cells, Respiration. 89 (2015) 572–582. https://doi.org/10.1159/000381905.

[37] J.D. Corrêa, C.M. Queiroz-Junior, J.E. Costa, A.L. Teixeira, T.A. Silva, Phenytoin-Induced Gingival Overgrowth: A Review of the Molecular, Immune, and Inflammatory Features, ISRN Dent. 2011 (2011) 1–8. https://doi.org/10.5402/2011/497850.

[38] L.W. Pappalardo, O.A. Samad, J.A. Black, S.G. Waxman, Voltage-gated sodium channel Nav1.5 contributes to astrogliosis in an in vitro model of glial injury via reverse Na+/Ca2+ exchange, Glia. 62 (2014) 1162–1175. https://doi.org/10.1002/glia.22671.

[39] Y. Kato, N. Koide, T. Komatsu, G. Tumurkhuu, J. Dagvadorj, K. Kato, T. Yokochi, Metformin attenuates production of nitric oxide in response to lipopolysaccharide by inhibiting MyD88-independent pathway, Horm. Metab. Res. 42 (2010) 632–636. https://doi.org/10.1055/s-0030-1255033.

[40] B.J. Davis, Z. Xie, B. Viollet, M.H. Zou, Activation of the AMP-activated kinase by antidiabetes drug metformin stimulates nitric oxide synthesis in vivo by promoting the association of heat shock protein 90 and endothelial nitric oxide synthase, Diabetes. 55 (2006) 496–505. https://doi.org/10.2337/diabetes.55.02.06.db05-1064.

[41] Y.W. Kim, S.Y. Park, J.Y. Kim, J.Y. Huh, W.S. Jeon, C.J. Yoon, S.S. Yun, K.H. Moon, Metformin restores the penile expression of nitric oxide synthase in high-fat-fed obese rats, J. Androl. 28 (2007) 555–560. https://doi.org/10.2164/jandrol.106.001602.

[42] T.R. O’Hora, F. Markos, N.F. Wiernsperger, M.I.M. Noble, Metformin causes nitric oxide-mediated dilatation in a shorter time than insulin in the iliac artery of the anesthetized pig, J. Cardiovasc. Pharmacol. 59 (2012) 182–187. https://doi.org/10.1097/FJC.0b013e31823b4b94.

[43] U. Förstermann, W.C. Sessa, Nitric oxide synthases: Regulation and function, Eur. Heart J. 33 (2012) 829. https://doi.org/10.1093/eurheartj/ehr304.

[44] N. Virgili, J.F. Espinosa-Parrilla, P. Mancera, A. Pastén-Zamorano, J. Gimeno-Bayon, M.J. Rodríguez, N. Mahy, M. Pugliese, Oral administration of the KATPchannel opener diazoxide ameliorates disease progression in a murine model of multiple sclerosis, J. Neuroinflammation. 8 (2011) 149. https://doi.org/10.1186/1742-2094-8-149.

[45] M.L. Garcia, V.F. King, J.L. Shevell, R.S. Slaughter, G. Suarez-Kurt&, R.J. Winquistqll, G.J. Kaczorowskill, Amiloride Analogs Inhibit L-type Calcium Channels and Display Calcium Entry Blocker Activity*, 1990. https://doi.org/10.1016/S0021-9258(19)39660-7.

[46] J.Á. Monsalve-Naharro, E. Domingo-Chiva, S.G. Castillo, P. Cuesta-Montero, J.M. Jiménez-Vizuete, Inhaled nitric oxide in adult patients with acute respiratory distress syndrome, Farm. Hosp. 41 (2017) 292–312. https://doi.org/10.7399/fh.2017.41.2.10533.

[47] J.H. Wandall, Effects of omeprazole on neutrophil chemotaxis, super oxide production, degranulation, and translocation of cytochrome b-245, 1992.

[48] F. Khalfi, B. Gressier, T. Dine, C. Brunet, M. Luyckx, L. Ballester, M. Cazin, J.C. Cazin, Verapamil inhibits elastase release and superoxide anion production in human neutrophils, Biol. Pharm. Bull. 21 (1998) 109–112. https://doi.org/10.1248/bpb.21.109.

[49] M. Suzuki, S. Teramoto, T. Matsuse, E. Ohga, H. Katayama, Y. Fukuchi, Y. Ouchi, Inhibitory effect of ambroxol on superoxide anion production and generation by murine lung alveolar macrophages, J. Asthma. 35 (1998) 267–272. https://doi.org/10.3109/02770909809068217.

[50] Ł. Bułdak, K. Łabuzek, R.J. Bułdak, M. Kozłowski, G. MacHnik, S. Liber, D. Suchy, A. Duława-Bułdak, B. Okopień, Metformin affects macrophages’ phenotype and improves the activity of glutathione peroxidase, superoxide dismutase, catalase and decreases malondialdehyde concentration in a partially AMPK-independent manner in LPS-stimulated human monocytes/macrophages, Pharmacol. Reports. 66 (2014) 418–429. https://doi.org/10.1016/j.pharep.2013.11.008.

[51] M.W. Verghese, R.C. Boucher, Effects of ion composition and tonicity on human neutrophil antibacterial activity, Am. J. Respir. Cell Mol. Biol. 19 (1998) 920–928. https://doi.org/10.1165/ajrcmb.19.6.3290.

[52] Y. Groemping, K. Rittinger, Activation and assembly of the NADPH oxidase: A structural perspective, Biochem. J. 386 (2005) 401–416. https://doi.org/10.1042/BJ20041835.

[53] Y. Xia, Superoxide generation from nitric oxide synthases, Antioxidants Redox Signal. 9 (2007) 1773–1778. https://doi.org/10.1089/ars.2007.1733.

[54] J. Frostegård, Immunity, atherosclerosis and cardiovascular disease, BMC Med. 11 (2013) 117. https://doi.org/10.1186/1741-7015-11-117.

[55] S. Gupta, N. Salam, V. Srivastava, R. Singla, D. Behera, K.U. Khayyam, R. Korde, P. Malhotra, R. Saxena, K. Natarajan, Voltage gated calcium channels negatively regulate protective immunity to Mycobacterium tuberculosis, PLoS One. 4 (2009) e5305. https://doi.org/10.1371/journal.pone.0005305.

[56] G. Li, X.P. Qi, X.Y. Wu, F.K. Liu, Z. Xu, C. Chen, X.D. Yang, Z. Sun, J.S. Li, Verapamil modulates LPS-induced cytokine production via inhibition of NF-kappa B activation in the liver, Inflamm. Res. 55 (2006) 108–113. https://doi.org/10.1007/s00011-005-0060-y.

[57] S. Song, T. Cheng, [The effect of systemic and local irradiation on wound macrophages and the repair promoting action of phenytoin sodium]., Zhonghua Yi Xue Za Zhi. 77 (1997) 54–57.

[58] M.L. Jackson, K.A. Ruppert, D.J. Kota, K.S. Prabhakara, R.A. Hetz, B.M. Aertker, S. Bedi, R.W. Mays, S.D. Olson, C.S. Cox, Clinical parameters affecting multipotent adult progenitor cells in vitro, Heliyon. 5 (2019) e02532. https://doi.org/10.1016/j.heliyon.2019.e02532.

[59] M.J. Thomassen, L.T. Buhrow, M.J. Connors, F.T. Kaneko, S.C. Erzurum, M.S. Kavuru, Nitric Oxide Inhibits Inflammatory Cytokine Production by Human Alveolar Macrophages, Am. J. Respir. Cell Mol. Biol. 17 (1997) 279–283. https://doi.org/10.1165/ajrcmb.17.3.2998m.

