## Supplementary Figure 1 for "Ion transport modulators differentially modulate inflammatory responses in THP-1 derived macrophages"

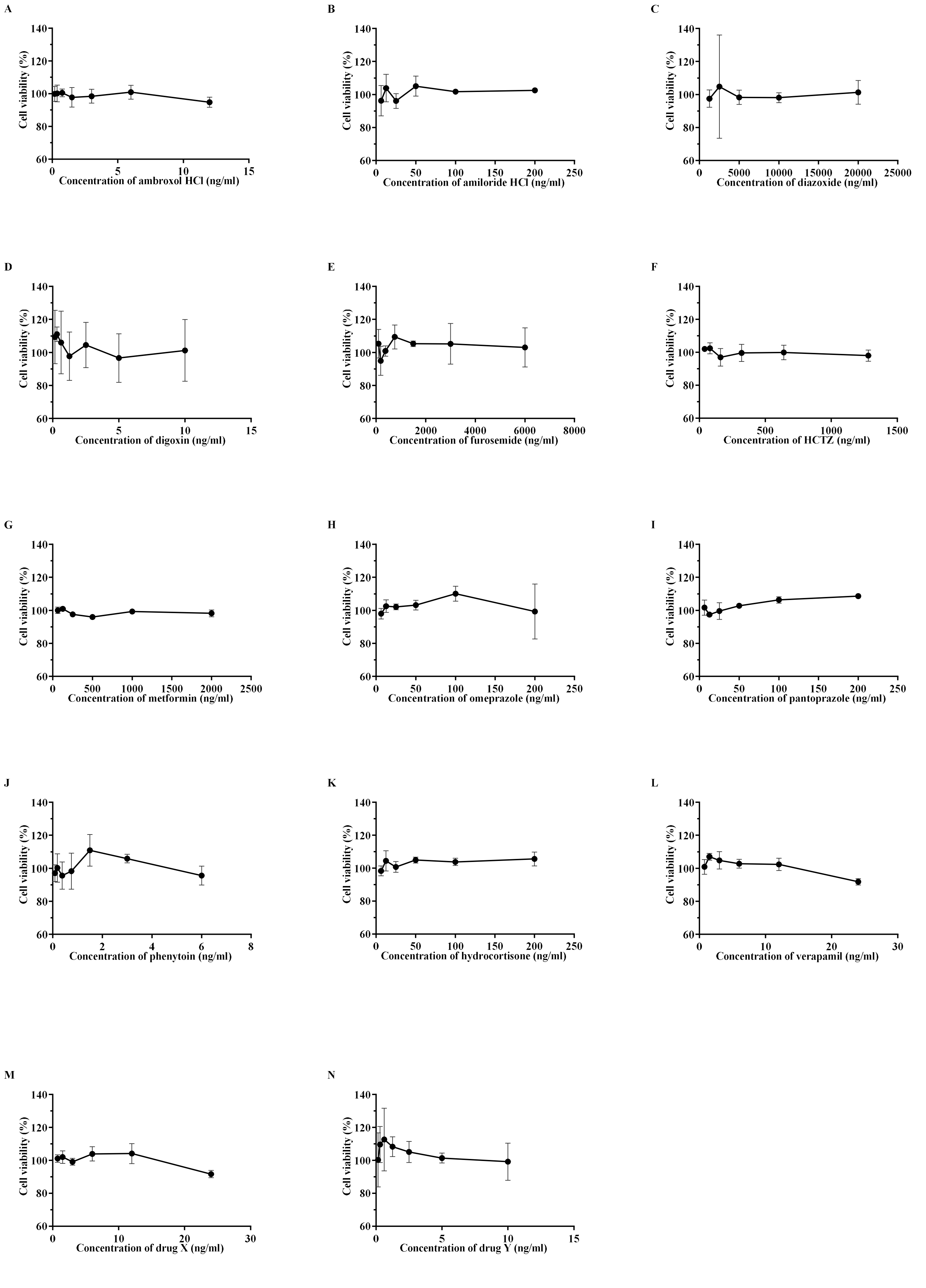


**Figure S1: Mean (+SD) viability of THP-1 derived macrophages following exposure to different concentrations of test drugs. HCTZ, hydrochlorothiazide. Reprinted from "Ion transport modulators as antimycobacterial agents," by SC Mitini-Nkhoma et al, 2020, Tuberculosis Research and Treatment, vol. 2020, https://doi.org/10.1155/2020/3767915.**
